# Phosphorylation of myosin A regulates *Plasmodium* sporozoite motility and is essential for efficient malaria transmission

**DOI:** 10.1101/2021.03.29.437488

**Authors:** Johanna Ripp, Xanthoula Smyrnakou, Marie-Therese Neuhoff, Friedrich Frischknecht

## Abstract

Malaria-causing parasites rely on an actin-myosin based motor for the invasion of different host cells as well as tissue traversal in mosquitoes and vertebrates. The unusual myosin A of *Plasmodium* spp. has a unique N-terminal extension which is important for red blood cell invasion by *P. falciparum* merozoites *in vitro* and harbors a phosphorylation site at serine 19. Here, using the rodent-infecting *P. berghei* we show that serine 19 is essential for efficient transmission of *Plasmodium* by mosquitoes as S19A mutants show defects in mosquito salivary gland entry and migration of salivary gland sporozoites in both 2D and 3D environments. Our data suggests that entry into salivary glands represents the strongest barrier in parasite transmission and hence is the key determinant for evolution of the motility and invasion machinery of these parasites.

**Highlights:** The unusual N-terminal extension of *Plasmodium* myosin A is important for efficient gliding motility

Altering the kinetics of the myosin A power stroke impacts *Plasmodium* life cycle progression and sporozoite motility

Myosin A phosphorylation at serine 19 is important for malaria transmission by mosquitoes

Salivary gland invasion emerges as key selection step for evolution of the parasite motor

## Introduction

Apicomplexan parasites rely on an actin-myosin-based motor for migration on and through tissues and for cell invasion (Heintzelman, 2015; Frénal *et al*, 2017). Compared with canonical actin-myosin motors, however, apicomplexans have evolved an intriguingly divergent machinery reliant on short and highly dynamic actin filaments and unique class XIV myosins, which lack the extended cargo-binding tail domain (Herm-Götz *et al*, 2002; Bookwalter *et al*, 2017; Douglas *et al*, 2018; Robert-Paganin *et al*, 2019). As key parts of the motor, actin 1 and myosin A (MyoA) are essential in *Plasmodium* spp., the causative agents of malaria in vertebrates (Bushell *et al*, 2017). Malaria is caused by the exponential replication of parasites in red blood cells. Red blood cells are invaded by otherwise largely non-migrating extracellular *Plasmodium* merozoites in an actin-myosin-dependent way (Mizuno *et al*, 2002; Das *et al*, 2017; Robert-Paganin *et al*, 2019; Blake *et al*, 2020). Mammal-infecting *Plasmodium* parasites undergo a complex life cycle that relies on *Anopheles* mosquitoes as disease vectors (Douglas *et al*, 2015). Indeed, *Plasmodium* parasites likely evolved in insects before they spread to vertebrate hosts suggesting that the core machinery of the parasite has initially adapted to a life in insects (Poinar, 2016). Within the mosquito, two highly motile extracellular forms of the parasite also rely on the actin-myosin motor. In the mosquito gut, ookinetes are formed and transmigrate across the midgut epithelium to transform into oocysts. In those oocysts, sporozoites are formed (Angrisano *et al*, 2012b). Sporozoites need motility to egress from oocysts, enter salivary glands, migrate in the skin, enter and exit the bloodstream and to infect hepatocytes (**Fig 1A**). This formidable journey likely requires a higher level of regulation of the actin-myosin motor compared to merozoites. Indeed, several actin regulatory proteins are only found to be important in sporozoites and mutations in actin or actin binding proteins could be identified that only impact sporozoite motility with little impact on merozoites or ookinetes (Ganter *et al*, 2009; Bane *et al*, 2016; Moreau *et al*, 2017; Douglas *et al*, 2018). Furthermore, MyoA and other components of the gliding machinery are phosphorylated in blood stages, ookinetes and sporozoites, suggesting that phosphorylation regulates motility (Sebastian *et al*, 2012; Alam *et al*, 2015; Lasonder *et al*, 2015; Swearingen *et al*, 2017).

**Figure 1.**
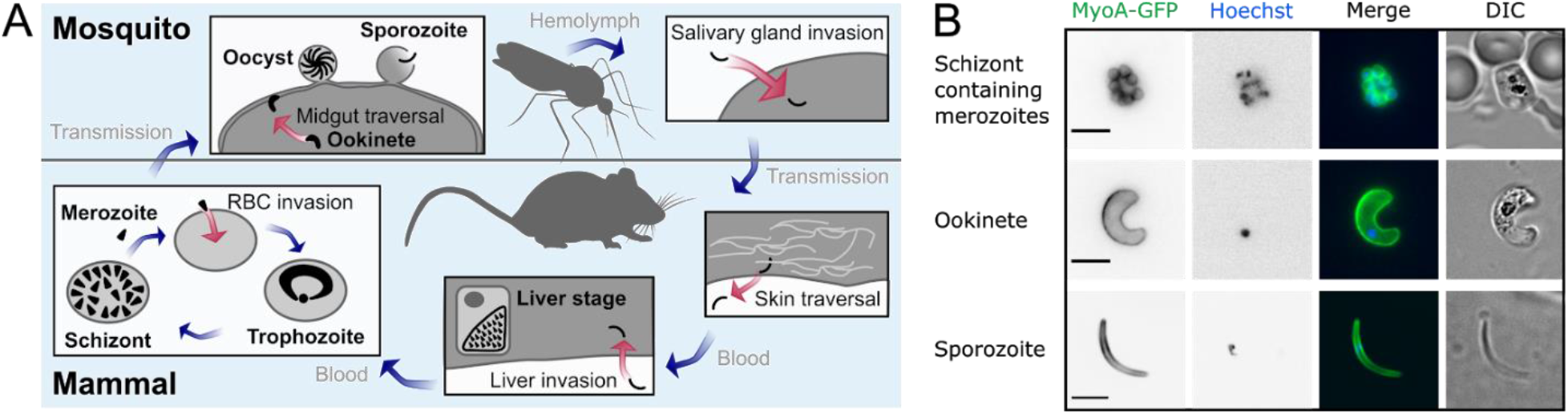
The myosin motor is required at multiple stages throughout the *Plasmodium* life cycle. (A) When an *Anopheles* mosquito feeds from an infected mammal, it takes up parasites with the bloodmeal. The parasites transform into motile ookinetes in the midgut. Myosin is essential for traversal of the midgut epithelium. Ookinetes transform into oocysts and develop into hundreds of sporozoites. Sporozoites egress from oocysts to float in the hemolymph. Active motility allows sporozoites to invade the salivary glands from where they are transmitted into the skin with the next bite. They traverse the skin and enter into a blood vessel. With the bloodstream they are transported until they reach the liver and actively invade a hepatocyte to develop into thousands of merozoites. These merozoites are released back into the bloodstream where they depend on myosin to actively invade red blood cells (RBCs) and multiply. Red and blue arrows indicate active and passive movements of the parasite, respectively. (B) The motor protein MyoA localizes to the periphery of motile *Plasmodium* stages. Images show parasites expressing GFP-tagged MyoA. Nuclei were stained with Hoechst. Scale bar, 5 µm.

Many insights into the function of the apicomplexan actin-myosin motor and its associated proteins have been gained from the study of *Toxoplasma gondii*, a highly successful parasite circulating between cats and their prey but also infecting about one third of the world’s human population (Frénal *et al*, 2017). For practical reasons, most studies on *T. gondii* are limited to tissue culture migration and invasion assays and it is important to note that the common ancestor of *T. gondii* and *Plasmodium* spp. split about 350-820 Mio years ago (Sato, 2011). Yet, both parasites, as well as most apicomplexans, feature similarly designed highly polarized invasive forms featuring a namesake complex apical end where vesicles fuse with the plasma membrane to secrete proteins or deposit them into the plasma membrane (Frénal *et al*, 2017). The plasma membrane is subtended by a membrane organelle called the inner membrane complex (IMC), which is the defining structure of the alveolates, the superphylum of organisms the apicomplexans belong to. On the cytoplasmic face of the IMC, a complex and stable membrane-associated network, the subpellicular network, is giving the parasites their shape (Harding & Frischknecht, 2020). In the narrow space (30 nm) between plasma membrane and IMC, the actin-myosin motor is located (Heintzelman, 2015; Frénal *et al*, 2017). Actin filaments are likely polymerized by formins located at the apical end of the parasites (Baum *et al*, 2008; Douglas *et al*, 2018; Tosetti *et al*, 2019). MyoA is anchored into the IMC by light chains and so-called gliding associated proteins (GAPs) and can move actin filaments from the front to the rear (Heintzelman, 2015; Frénal *et al*, 2017). As the actin filaments bind receptors spanning the plasma membrane these receptors are moved rearwards. If the receptors engage with ligands on the surface of a cell, tissue or glass coverslip, the parasite moves forward. While actin filaments could not yet be clearly shown between IMC and plasma membrane in extracellular *Plasmodium* stages (Kudryashev *et al*, 2010; Angrisano *et al*, 2012a; Siden-Kiamos *et al*, 2012), MyoA has been localized to the IMC of all invasive stages through expression of a GFP-fusion protein (Wall *et al*, 2019) (**Fig 1B**).

Conditional deletion of *myoA* or *actin* in *T. gondii* showed a dramatic decrease in both tachyzoite motility and cell invasion (Meissner *et al*, 2002; Andenmatten *et al*, 2013; Egarter *et al*, 2014). Replacing the *myoA* promoter with a promoter that is only active in blood stages in the rodent-infecting malaria parasite *P. berghei* showed that ookinetes are still formed but cannot migrate in the absence of MyoA (Siden-Kiamos *et al*, 2011). Conditional mutations in MyoA and a myosin light chain in *P. falciparum* revealed an orchestrated need for merozoite force production to enter red blood cells (Blake *et al*, 2020). Crystal structures of *P. falciparum* MyoA showed that interaction sites formed by E6 and phosphorylated S19 in the N-terminal extension are important for MyoA kinetics (Robert-Paganin *et al*, 2019; Moussaoui *et al*, 2020). Mutations of these amino acids leading to the disruption of the respective interactions reduced the speed at which purified MyoA transports actin filaments *in vitro* but increased maximal force production (Robert-Paganin *et al*, 2019; Moussaoui *et al*, 2020). This suggests that phosphorylation of S19 is required for maximum myosin velocity and might hence play a role in ookinetes or sporozoites, that rely on motility for much longer periods than merozoites. In laboratory settings, the major human infecting *P. falciparum* is usually studied in cultured blood cells, while for the study of mosquito transmission the rodent malaria parasite *P. berghei* is used for reasons of experimental ease, ethical limitations and safety (Matz & Kooij, 2015). In order to probe the role of the N-terminal extension of MyoA for ookinete and sporozoite motility, we sought to generate parasite lines of *P. berghei* that harbour different mutations in MyoA and analyse their effects in the mosquito stages. We show that the endogenous 3’ untranslated region (UTR) of *myoA* is important for expression in sporozoites limiting gene modifications by classic homologous recombination. Consistent with data from *P. falciparum*, we find that the N-terminal extension is essential for blood stages. However, point mutations of E6 and S19 yielded viable parasite lines that can infect mosquitoes at normal levels. Strikingly, parasites expressing S19A but not the phosphomimetic S19D showed reduced mosquito salivary gland colonization and liver infection suggesting that phosphorylation of S19 is essential for efficient transmission.

## Results

### Exchange of 3’myoA by 3’dhfs leads to reduced myoA expression levels resulting in a defect in salivary gland invasion

To understand how single amino acid residues contribute to the function of MyoA, we first attempted to use conventional gene editing of the *myoA* locus which results in integration of the 3’UTR of *dhfs* and a selection cassette downstream of *myoA* (**Supplementary Fig 1**). This would allow insertions of mutations similar to what was achieved in our study on the actin binding proteins coronin and profilin (Bane *et al*, 2016; Moreau *et al*, 2017). Along with the generation of the control line, we also tested whether the *P. falciparum* orthologue can complement the function of *P. berghei* MyoA (PbMyoA) by replacing PbMyoA with *P. falciparum myoA* (PfMyoA). In addition, we C-terminally tagged the protein with GFP for localization studies (**Fig 2A and Supplementary Fig 1**). To test if the clonal lines proceed through the life cycle, a prerequisite for the mutagenesis study, we analyzed all life cycle stages. While the blood stage growth rate of the PbMyoA 3’dhfs and PbMyoA-GFP 3’dhfs parasite lines were similar to wild type (wt), it was significantly reduced from 9-fold per day in wt to 6-fold per day in the PfMyoA 3’dhfs line (**Fig 2B**). To assess the ability of parasites to infect mosquitoes, we counted the numbers of oocysts that formed in the mosquito midguts 10 to 14 days after the insects took a blood meal on infected mice. All parasite lines could form oocysts, but the PbMyoA 3’dhfs and PfMyoA 3’dhfs lines showed significantly reduced numbers of oocysts (**Fig 2C**). In line with the very low number of oocysts, the number of sporozoites that formed in midgut oocysts was most reduced in the PfMyoA 3’dhfs line (**Table 1**). In contrast, PbMyoA 3’dhfs and PbMyoA-GFP 3’dhfs sporozoite numbers within oocysts were similar to wt. However, sporozoite numbers from salivary glands were strongly reduced, suggesting a defect in salivary gland invasion (**Table 1, Fig 2D**). Surprisingly, the ratio of salivary gland to midgut sporozoites was similar to wt in the PfMyoA 3’dhfs parasite line. Next, we examined whether the defect in salivary gland invasion could be due to reduced motility of sporozoites floating in the hemolymph, the circulating fluid of the mosquito. Indeed, a much lower fraction of isolated hemolymph sporozoites displayed gliding motility on glass in these parasite lines (**Fig 2E**). We hypothesized that the defect in *in vitro* motility and salivary gland invasion might be due to a difference in *myoA* expression resulting from the replaced 3’UTR. Indeed, quantification of *myoA* mRNA levels via qPCR revealed a 5 to 10-fold reduction in the PbMyoA 3’dhfs line in early oocysts, while there was increased *myoA* expression in schizonts (**Fig 2F**). Hence a different strategy for mutagenesis was needed that kept the endogenous 3’UTR of *myoA* intact.

**Figure 2.**
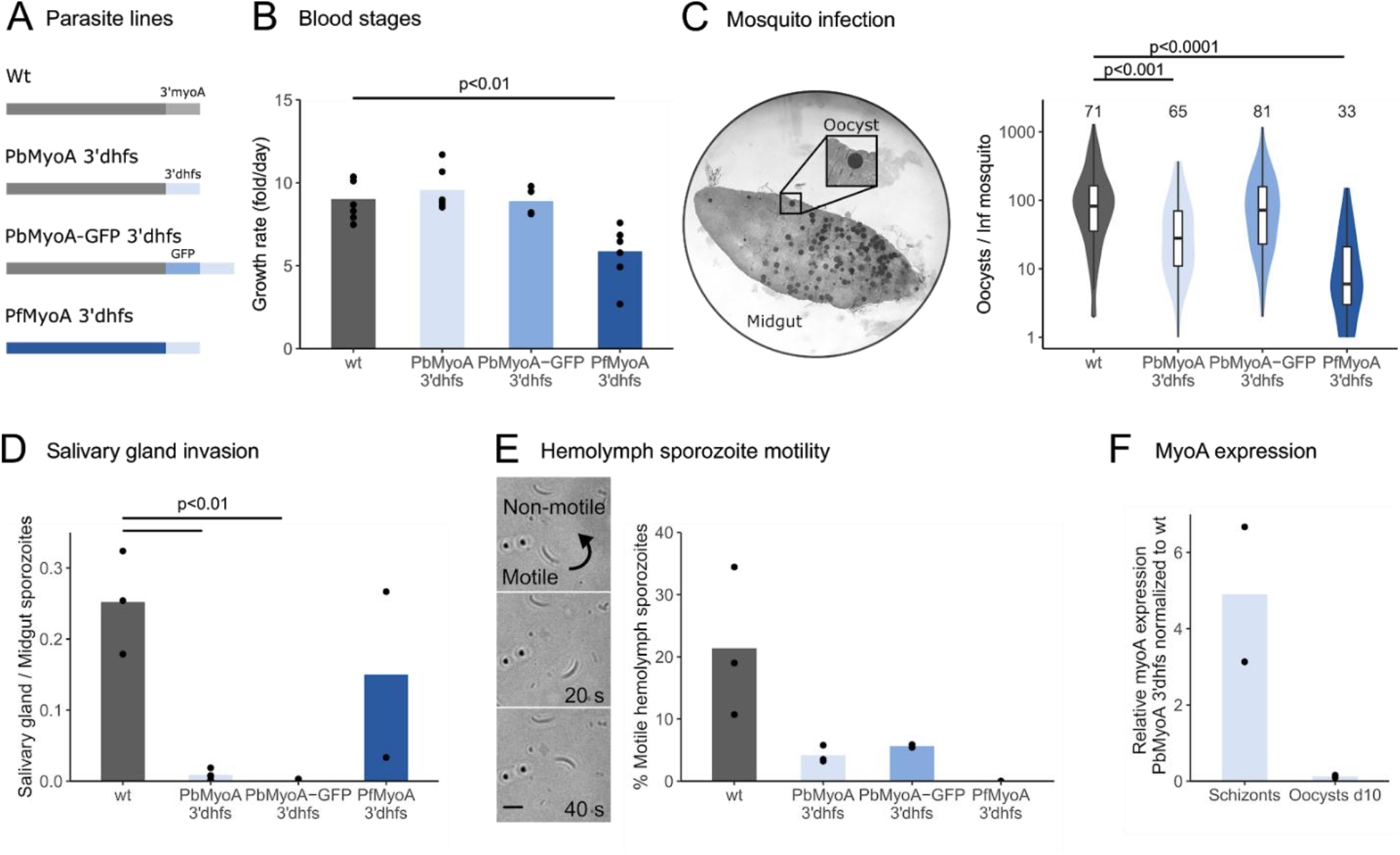
Effect of 3’UTR replacement of *myoA* by the 3’UTR of *dhfs* on life cycle progression and *myoA* expression. (A) Scheme showing simplified *myoA* loci in transgenic parasite lines. (B) Blood stage growth rate of mutated parasite lines. A single parasite was injected *i*.*v*. into a mouse and the growth rate was calculated from parasitemia at day 6-9 for each mouse (black dots). Bars represent the mean of 4-7 mice. (C) Infected midgut showing oocysts stained with mercurochrome. The graph shows oocyst numbers per infected mosquito. Numbers of mosquitoes that were analyzed are indicated above the graph. Data derived from at least two independent cage feeds. Box and whisker plots depict the 25% quantile, median, 75% quantile, and nearest observations within 1.5 times the interquartile range (whiskers). (D) Ratio of salivary gland to midgut sporozoites of the indicated parasite lines. Black dots: individual experiments with 10-20 mosquitoes. Bars: mean. (E) Hemolymph sporozoites were isolated and observed on glass slides by live cell imaging. Arrow indicates direction of a motile sporozoite. The graph shows the fraction of motile hemolymph sporozoites. Black dots: individual experiments with 10-20 mosquitoes. Bars: mean. (F) qRT–PCR analysis of schizonts and oocysts of the PbMyoA 3’dhfs parasite line. Individual data points correspond to the mean of three technical replicates. Significance for (B), (D) and (E) determined by One-way analysis of variance with Tukeys Multiple Comparison test. Significance for (C) determined by Kruskal-Wallis test with Bonferroni’s Multiple Comparison test.

**Table 1.** Mosquito infection capacity of the different parasite lines

| Parasite line | Infection rate [%] (n) | MG sporozoites/<br>inf mosquito (n) | HL sporozoites/<br>inf mosquito (n) | SG sporozoites/<br>inf mosquito (n) |
| --- | --- | --- | --- | --- |
| wt | 80 ± 10 (85) | 76000 ± 42000 (90) | 3000 ± 100 (37) | 20000 ± 14000 (50) |
| PbMyoA 3'dhfs | 60 ± 20 (127) | 83000 ± 34000 (123) | 6300 ± 3400 (61) | 700 ± 800 (59) |
| PbMyoA-GFP 3'dhfs | 80 ± 20 (115) | 89000 ± 57000 (59) | n.d. | **100 ± 100** (57) |
| PfMyoA 3'dhfs | 50 ± 20 (73) | **7500 ± 6400** (60) | n.d. | **600 ± 300** (56) |
| RL | 0 (40) | n.d. | n.d. | n.d. |
| wt recon | 70 ± 20 (91) | 84000 ± 36000 (74) | 5600 ± 1800 (56) | 13000 ± 6900 (48) |
| MyoA E6R | 80 ± 20 (112) | 70000 ± 15000 (99) | 4700 ± 1000 (51) | 16000 ± 7200 (48) |
| MyoA S19A | 60 ± 20 (117) | 130000 ± 70000 (87) | 11000 ± 6900 (38) | **3900 ± 2600** (48) |
| MyoA S19D | 80 ± 10 (90) | 78000 ± 50000 (104) | 4300 ± 1800 (49) | 10000 ± 2600 (54) |

Infection rate: numbers of infected mosquito after a cage feed. n: numbers of mosquitoes investigated.
Note that sporozoite numbers are from at least two different cage feeds determined from pooled organs. Significant reductions are shown in bold.

### Phosphorylation of MyoA at serine 19 is important for salivary gland invasion of sporozoites and efficient malaria transmission

In order to leave the 3’UTR within the myoA locus intact, we next used a cloning strategy that resulted in selection marker free parasite lines (Lin et al., 2011). In a first step, we generated a clonal ama1 promoter swap parasite line (**Supplementary Fig 2**). In a second step, we replaced the ama1 promoter by the endogenous myoA promoter and the *myoA* ORF carrying the mutation of interest at the 5’ end of *myoA*. As a control, we reconstituted the wt *myoA* locus. This resulted in control parasites proceeding through the life cycle as anticipated (**Table 1, Fig 3**). To assess the role of the unique N-terminus of MyoA on parasite motility, we focused on two amino acids, E6 and S19, which are thought to be especially important as they interact with switch I/II and the converter, respectively and are required to transport actin filaments at maximum speed (Robert-Paganin *et al*, 2019; Moussaoui *et al*, 2020) (**Fig 3A**). To this end, we introduced a reverse charge at position 6 (mutant E6R) and replaced serine at position 19 with either a phosphorylation incompatible alanine (mutant S19A) or a phosphomimetic aspartate (mutant S19D) (**Fig 3B**). Previous data suggests that the S19A mutant would only have a minor impact on merozoites but affect the fast gliding stages (Robert-Paganin *et al*, 2019; Blake *et al*, 2020). To study the overall importance of the N-terminal extension, we also generated a construct aiming to delete the first 19 amino acids from the N-terminus of MyoA (**Fig 3B**). While we obtained parasites with the point mutations after transfection, we failed to delete the N-terminal extension in four attempts, suggesting that this part of the protein is essential. Interestingly, E6R and S19A mutants grew as well as wild type while S19D parasites grew significantly slower in the blood (**Fig 3C**). In contrast, mutagenesis had no effect on mosquito infection as assessed by the numbers of oocysts present in mosquitoes fed on infected mice (**Fig 3D**). This indicates that the mutations had no major effect on ookinete formation and migration. Oocysts produced normal numbers of sporozoites in all lines, which were able to exit into the hemolymph. However, sporozoite numbers in the hemolymph were slightly enhanced in the S19A parasite line, resulting in an increase in the ratio of hemolymph to midgut sporozoites (**Table 1, Fig 3E**). Far fewer sporozoites were found in salivary glands of this S19A parasite line than in the other lines (**Table 1**) as evidenced by a significantly lower ratio of salivary gland to midgut sporozoites (**Fig 3F**). This suggests a defect of S19A sporozoites to enter into salivary glands.

**Figure 3.**
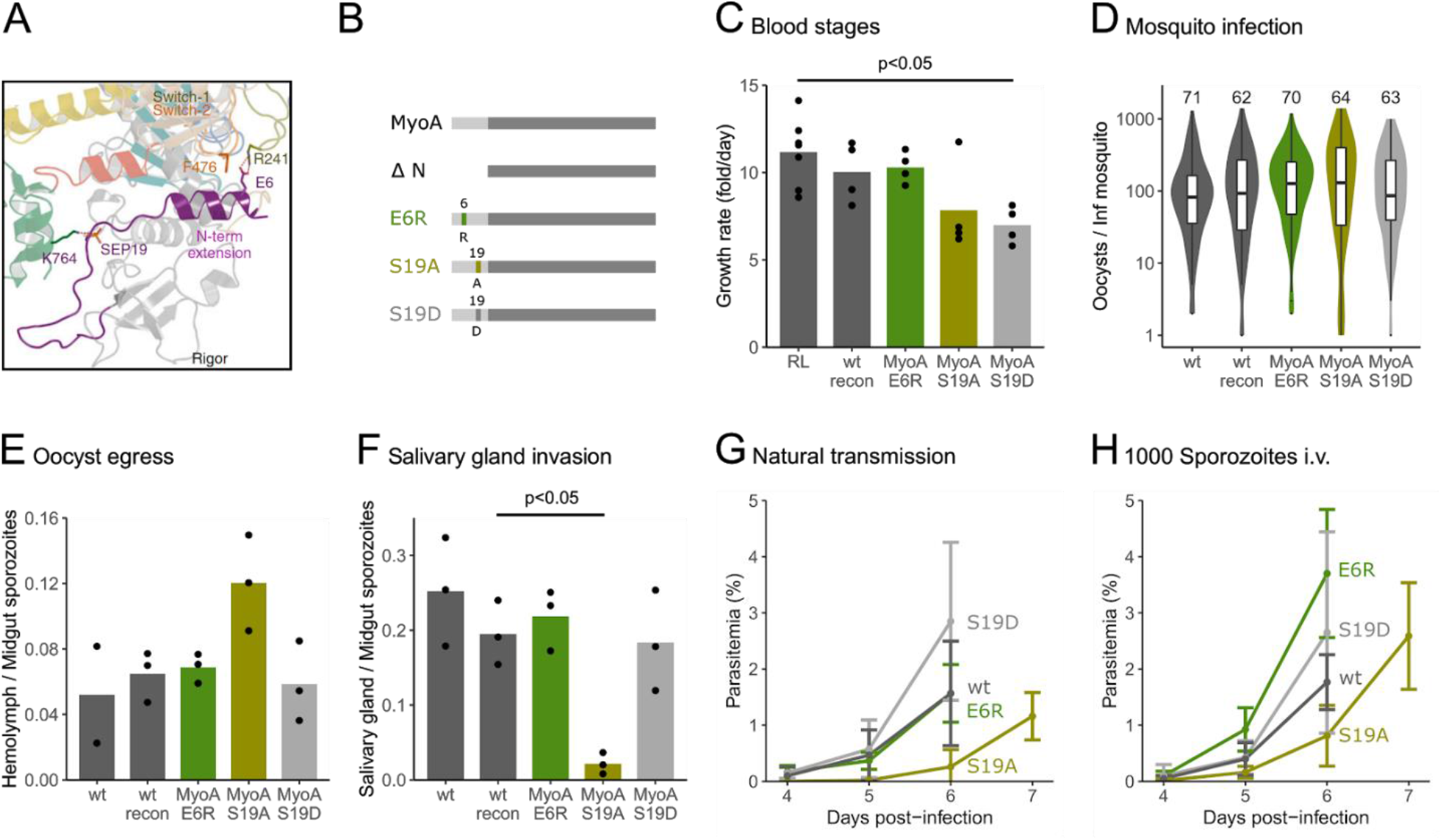
Phosphorylation of MyoA at serine 19 is important for salivary gland invasion of sporozoites. (A) Zoom of the PfMyoA structure showing important amino acid interactions for stabilization of the rigor-like state that are thought to influence the kinetic properties of MyoA. The N-terminal extension is depicted in purple. It is located in proximity to Switch I (green) and the connectors Switch II (orange) and Relay (yellow). Phosphorylated serine 19 (SEP19) in the N-terminal extension interacts with lysine 764 (K764) in the converter and glutamic acid 6 (E6) in the N-terminal extension interacts with arginine 241 (R241) as well as phenylalanine 476 (F476) from switch I and II. Image taken from Robert-Paganin et al. (2019). The image was published under a Creative Commons Attribution 4.0 International License (http://creativecommons.org/licenses/by/4.0/). (B) Scheme of mutated MyoA versions that are expressed in clonal parasite lines at the endogenous *myoA* locus. Either the complete N-terminal extension is depleted (19 amino acids) or point mutations within amino acids that are located in the N-terminal extension are introduced. (C) Blood stage growth rate of clonal parasite lines. A single parasite or 100 iRBCs were injected *i*.*v*. into a mouse and the growth rate was calculated from parasitemia at day 6-9. Black dots represent a single mouse; bars represent the mean. (D) Number of oocysts per infected mosquito. Numbers of analyzed mosquitoes are shown above the graph. Data from at least three independent cage feeds. Box and whisker plots depict the 25% quantile, median, 75% quantile, and nearest observations within 1.5 times the interquartile range (whiskers). (E) Ratio of hemolymph to midgut sporozoites. Black dots: individual experiments with 10-20 mosquitoes. Bars: mean. (F) Ratio of salivary gland to midgut sporozoites. Black dots: individual experiments with 10-20 mosquitoes. Bars: mean. Significance for (D) determined by Kruskal-Wallis test with Bonferroni’s Multiple Comparison test. Significance for (C), (E) and (F) determined by One-way analysis of variance with Tukeys Multiple Comparison test. (G) Blood stage infection of mice after transmission by mosquito bite. Shown is the mean ± standard deviation. (H) Blood stage infection of mice after *i*.*v*. injection of 1000 sporozoites.

Next, we tested whether the mutations within the N-terminal extension of MyoA have an effect on parasite transmission back to mice. To this end, we allowed infected mosquitoes to feed on mice. We then determined the appearance and numbers of infected red blood cells starting three days after infection to evaluate the success of parasite transmission. The S19A parasite line only infected 60% of mice (5 out of 8) via natural transmission by mosquito bite, while all other parasite lines infected 100% of mice (**Table 2**). Those mice that developed a blood stage infection with the S19A mutant showed a one-day delay in infection as compared to the wt control (**Table 2, Fig 3G**). A one-day delay without decreased numbers of infected mice is generally considered a 90% reduction of infectivity (Vanderberg, 1975). These findings suggest that phosphorylation at S19 is required for efficient salivary gland invasion and efficient malaria transmission. Such a defect in natural transmission can be due to reduced numbers of sporozoites in the salivary glands, which leads to fewer transmitted sporozoites (Aleshnick *et al*, 2020), a defect in motility in the skin or a defect in liver cell invasion (Frischknecht & Matuschewski, 2017). To test for a role in transmission and transmigration of the skin versus liver cell invasion, we intravenously injected 1000 salivary gland derived sporozoites. In this experiment all mice, including those infected with S19A sporozoites, became infected. However, there was still a delay in S19A infection (**Table 2, Fig 3H**). While not excluding that S19A mutants have a defect in skin migration, this indicates that S19A parasites have a reduced capacity to invade the liver.

**Table 2.** Transmission capacity of the different parasite lines

| Parasite line | Natural transmission (10 mosquito bites) |  |  | 1000 sporozoites i.v. |  |  |
| --- | --- | --- | --- | --- | --- | --- |
|  | Prepatency | Parasitemia d6 | Infected mice | Prepatency | Parasitemia d6 | Infected mice |
| wt recon | 3.8 ± 0.8 | 1.6 ± 0.9 | 6/6 | 3.5 ± 0.5 | 1.8 ± 0.5 | 6/6 |
| MyoA E6R | 4.0 ± 0.0 | 1.6 ± 0.5 | 3/3 | 3.3 ± 0.5 | 3.7 ± 1.1 | 4/4 |
| MyoA S19A | **5.0 ± 0.0** | **0.3 ± 0.3** | **5/8** | **4.1 ± 0.4** | **0.8 ± 0.5** | 8/8 |
| MyoA S19D | 3.3 ± 0.5 | 2.9 ± 1.4 | 4/4 | 3.8 ± 0.5 | 2.7 ± 1.8 | 4/4 |

Prepatency: day after mouse infection where parasites appear for the first time in a blood smear.
Significant reductions are shown in bold.

### Mutated MyoA affects sporozoite motility

*Plasmodium* sporozoites can migrate at more than 1 µm/s, an order of magnitude faster than neutrophils (Vanderberg, 1974). Some even reach peak instantaneous velocities of over 5 µm/s (Münter *et al*, 2009). To investigate whether the defect of S19A parasites in salivary gland infection can be explained by reduced motility, we probed hemolymph and salivary gland sporozoite motility of all generated parasite lines on glass. This showed that around 20% of wt hemolymph sporozoites could migrate in the typical circular path of sporozoites or in a back-and-forth manner (termed patch gliding) while a reduced fraction of just 3% of S19A hemolymph sporozoites were motile (**Fig 4A**). The S19D mutant also showed a slight reduction of motile hemolymph sporozoites. We determined the speed of the migrating sporozoites isolated from the hemolymph and found that both S19A and S19D hemolymph sporozoites migrated significantly slower as compared to the reconstituted wt sporozoites. We next isolated sporozoites from salivary glands and imaged their migration behavior. Here, the fraction of motile E6R and S19A salivary gland sporozoites was slightly lower but not significantly reduced compared to wt. However, the speed of all mutant parasites was significantly reduced as compared to wt sporozoites (**Fig 4B**). As some sporozoites expressing mutated actin-binding proteins alter their migration paths (Montagna *et al*, 2012; Bane *et al*, 2016), we next compared the trajectories of the migrating sporozoites from the different parasite lines. This showed no apparent difference between the mutant hemolymph and salivary gland sporozoites and the wt control parasites (**Fig 4C, D**).

**Figure 4.**
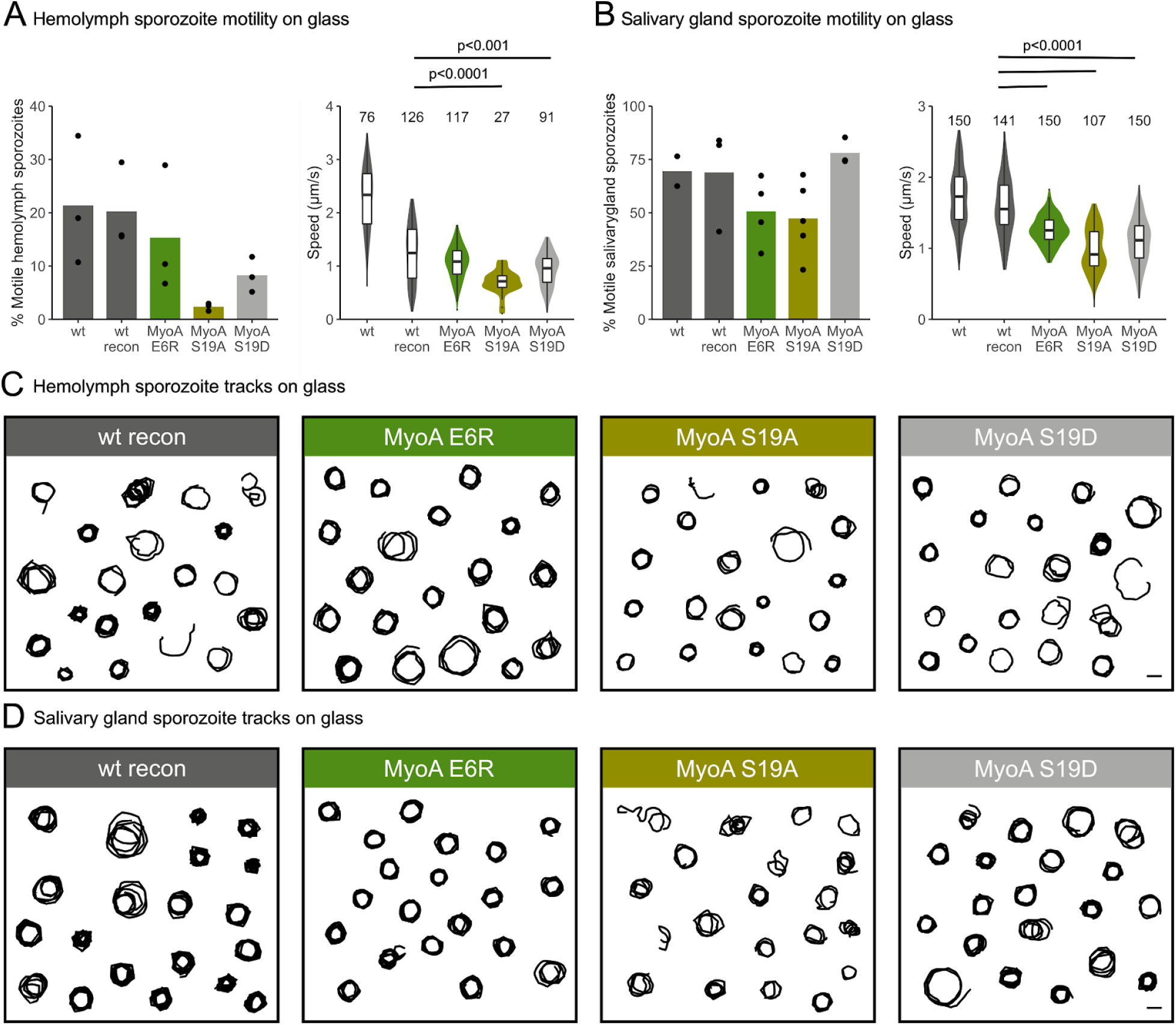
Reduced speed during *in vitro* sporozoite gliding motility of parasite lines expressing mutated MyoA. (A) Fraction (left) and speed (right) of motile hemolymph sporozoites on glass. (B) Fraction (left) and speed (right) of motile salivary gland sporozoites on glass. Dots correspond to individual experiments and bars indicate the mean. Box and whisker plots depict the 25% quantile, median, 75% quantile, and nearest observations within 1.5 times the interquartile range (whiskers). Significance for (A left) and (B left) determined by One-way analysis of variance with Tukeys Multiple Comparison test. Significance for (A right) and (B right) determined by Kruskal-Wallis test with Bonferroni’s Multiple Comparison test. Selected trajectories of 20 manually tracked sporozoites from hemolymph (C) and salivary glands (D) moving over a period of 3 min. Scale bar, 10 μm.

### Changing MyoA kinetics affects sporozoite force production

Parasite motility results from a complex interplay between adhesion and force generation. Yet, force generation of sporozoites has so far only been investigated in parasites lacking either surface receptors or actin-binding proteins without a direct function in force generation (Münter *et al*, 2009; Hegge *et al*, 2012; Quadt *et al*, 2016; Moreau *et al*, 2017, 2020). Intriguingly, force generation of sporozoites was found to be lower in mutant parasites that show similar migratory capacity or speeds as wt. To test whether mutations that are thought to impact MyoA kinetics and force generation on the molecular level result in a difference in sporozoite force production on the cellular level, we used a previously established assay based on optical tweezers (Quadt *et al*, 2016). To this end, a polystyrene bead was kept in the laser trap at a constant trapping force and positioned at the apical end of a moving salivary gland derived sporozoite. We then investigated whether a sporozoite could pull the bead out of the trap. As we used a new commercial setup instead of the one described before (Quadt *et al*, 2016), we first calibrated the tweezers. After calibration, we repeatedly observed that around 70% of wt sporozoites were able to pull a bead out of the trap at 30 pN of force. This percentage was observed at a force of 70 pN in the previous setup (Quadt *et al*, 2016; Moreau *et al*, 2017, 2020). Currently we do not know the source of this difference and as the previous setup was dismantled, we unfortunately cannot compare them side-by-side. Nevertheless, while about 70% of wt or wt-like sporozoites were able to pull the bead out of the trap, only 18% of the E6R mutant and 25% of the S19D mutant were capable of pulling the bead out of the trap (**Fig 5A**). We were not able to perform these experiments with the phosphodeficient mutant due to the low number of S19A sporozoites in the mosquito salivary gland.

**Figure 5.**
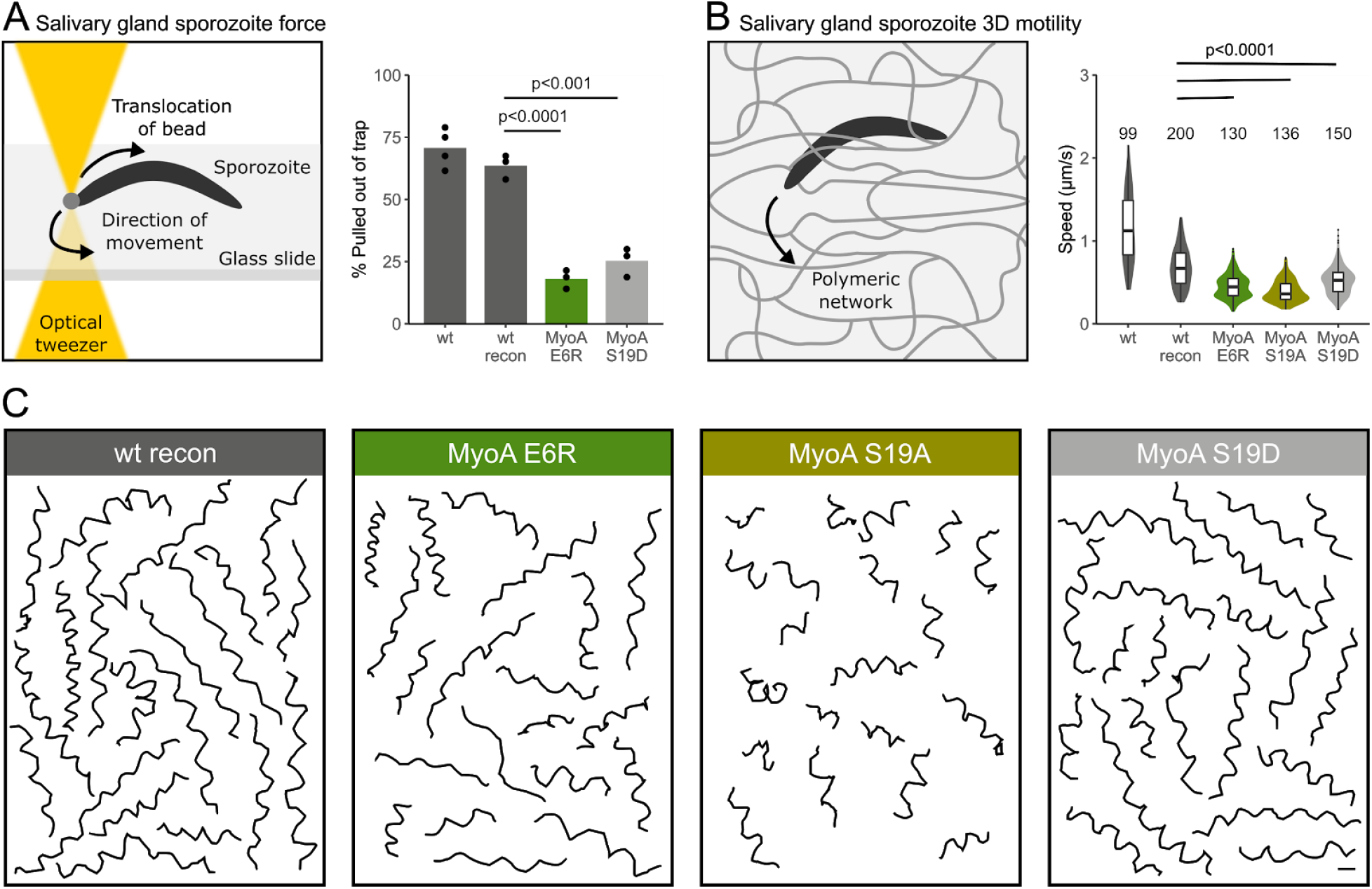
Lower force generation capacity and 3D motility of parasite lines expressing mutated MyoA. (A) Scheme depicting the setup of optical tweezers for measuring the forces that a sporozoite can generate to pull a bead out of an optical trap. The bead is placed at the apical end of the sporozoite. The graph shows the fraction of sporozoites that pulled the bead out of the optical trap. Black dots: individual experiments. Bars: mean. 73-131 sporozoites were probed for each line. Significance determined by One-way analysis of variance with Tukeys Multiple Comparison test. (B) Scheme showing a sporozoite moving through a polymeric network in 3D. The graph shows sporozoite speeds as measured by manual tracking from imaging of sporozoites in a 3D hydrogel. Box and whisker plots depict the 25% quantile, median, 75% quantile, and nearest observations within 1.5 times the interquartile range (whiskers). Significance determined by Kruskal-Wallis test with Bonferroni’s Multiple Comparison test. (C) Selected trajectories of 20 moving sporozoites observed over a period of 3 min. Scale bar, 10 μm.

### S19A mutation diminishes 3D migration of sporozoites

We next investigated 3D motility of the mutants in polyacrylamide-based hydrogels as a model to study their migration through the skin (Ripp *et al*, 2021). To this end, dissected salivary glands were sandwiched between two hydrogels and the sporozoites released from the gland entered into the gel on corkscrew-like paths. This allowed us to also image the S19A salivary gland derived sporozoites. Previous experiments revealed that mutant sporozoites with severe defects in 2D motility due to changes in their substrate adhesion capacity showed much improved migration capacity in 3D (Ripp *et al*, 2021). In contrast, the speed of all sporozoites expressing myosin mutants was significantly reduced compared to the wt-like control line (**Fig 5B**). The trajectories from sporozoites tracked for three minutes of observation were shorter, especially for the S19A mutant highlighting the lower displacement of these sporozoites (**Fig 5C**). This, along with the *in vivo* transmission data, is suggestive of a migration defect in the skin.

Taken together, these data suggest that ablating stabilizing interactions in the rigor state of MyoA results in a defect in force generation and motility at the sporozoite stage. While the E6R and S19D mutations only have a minor impact, the S19A mutation fails to efficiently enter into salivary glands. Entry into salivary glands might therefore constitute the strongest barrier in parasite transmission and hence represent the main obstacle against which the parasites needed to primarily evolve their migration and invasion machinery.

## Discussion

Malaria parasites use substrate-based and myosin-powered gliding motility to enter into red blood cells, traverse the midgut epithelium, enter into salivary glands, migrate in the skin, enter and exit blood vessels, and invade hepatocytes. While entry into red blood cells is completed in less than one minute, migration in the skin can last tens of minutes and ookinetes can migrate for several hours (Gilson & Crabb, 2009; Hopp *et al*, 2015; Moreau *et al*, 2017; Ripp *et al*, 2021; Hopp *et al*, 2021). Although invasion of the red cells appears as the easiest of these tasks for the parasite due to the short time and distance, recent work has shown that it presents an intriguing barrier with precise physical thresholds to be surmounted by the parasite (Blake *et al*, 2020; Kariuki *et al*, 2020). A malaria-protective polymorphism in the Dantu blood group was found to be based on an elevated mean membrane tension from 60 to 90 μN/m with 40 μN/m serving as a threshold above which no invasion can take place (Kariuki *et al*, 2020). This suggests that a large number of merozoites do not enter red cells due to this threshold. Likely, a similar phenomenon also applies for entry into salivary glands as in most mosquito infections a majority of parasites produced in the oocysts do not manage to enter into the glands. Generation of different *P. falciparum* parasite lines with mutated MyoA revealed the necessity for force generation by the merozoite in its step-wise process of red cell invasion (Blake *et al*, 2020). To complement the studies performed in *P. falciparum* and analyze the effect of MyoA mutations *in vivo* as well as throughout the whole life cycle, we generated *P. berghei* parasite lines harbouring mutations within the N-terminal extension of MyoA. Subtle effects as have been seen in *P. falciparum* invasion assays were not detected during the blood stage growth of our lines suggesting that they play comparatively minor roles *in vivo* in the rodent model. Similarly, we observed a measurable reduction in force generation by two mutants in *P. berghei* sporozoites. Yet this defect did not translate into a significant loss in mosquito-to-mouse transmission efficiency of the parasite lines. Only the S19A mutant line, which cannot be phosphorylated at S19 and hence alters the kinetics of the MyoA power stroke (Robert-Paganin *et al*, 2019) showed a dramatic reduction in the ability to transmit from mosquito to mammal (**Fig 6**). This block was mainly due to the incapacity of the parasites to colonize the salivary glands of the mosquito, although the parasite also showed defects in gliding in a skin-like gel and in liver infection. These results indicate that sporozoites depend on very fast myosin dynamics for efficient transmission.

**Figure 6.**
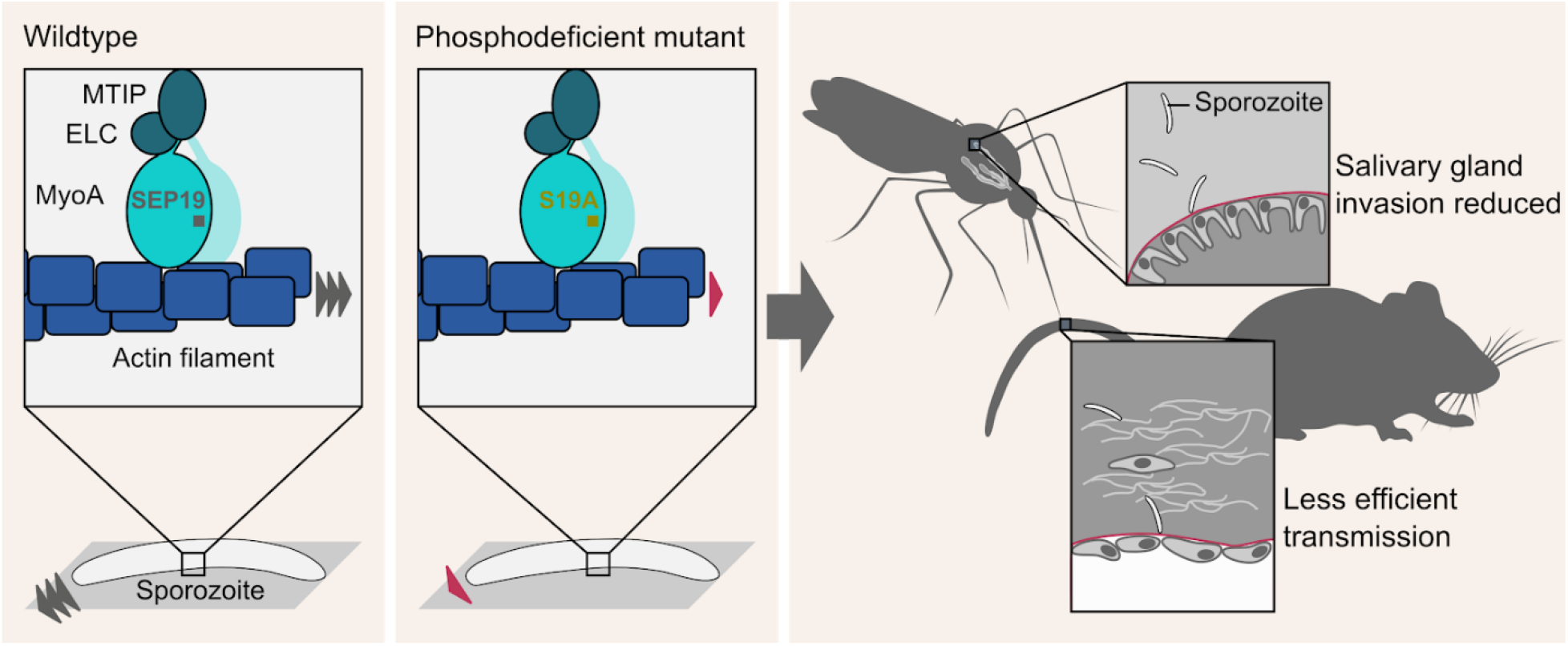
Cartoon portraying importance of MyoA phosphorylation during transmission of Plasmodium. Wildtype myosin is phosphorylated in early sporozoites allowing the generation of optimal force transduction and gliding motility. Dephosphorylation is also important for optimal force generation as S19D shows lower force and speed. Non-phosphorylated myosin still allows sporozoite gliding but at reduced speed with too little force generated for efficient salivary gland invasion. Disruption of the interaction of the N-terminus with the Switch I and II (E6R mutation) has similar impact as constitutive phosphorylation as mimicked by S19D.

Similar to our mutants, *T. gondii* MyoA mutants that could not be phosphorylated showed changes in their motile behavior and a decreased capacity to exit and enter cultured host cells (Tang *et al*, 2014). In *T. gondii*, phosphorylation of MyoA is dependent on calcium and the calcium dependent protein kinase TgCDPK3 was identified as a kinase to phosphorylate serines in the N-terminus of MyoA (Tang *et al*, 2014; Gaji *et al*, 2015). Calcium is also important for activation of *Plasmodium* sporozoite motility (Carey *et al*, 2014) and three CDPKs have been shown to impact sporozoite motility with two influenced by protein kinase G (Govindasamy *et al*, 2016; Govindasamy & Bhanot, 2020). Our data showing that S19 phosphorylation is important during salivary gland invasion, migration in the skin and liver invasion, suggests that similar signaling pathways play a role at different steps during malaria transmission.

Previous data from the mutagenesis of the ubiquitously expressed actin-binding protein profilin (Moreau *et al*, 2017, 2020) suggested a hierarchy of properties that are important for parasite transmission. The parasite can tolerate small perturbations that lead to a decrease in force generation without showing a defect in gliding motility and transmission capacity of the sporozoite (Moreau *et al*, 2017). Successively larger perturbations first lead to a drop in sporozoite gliding efficiency followed by a reduction in salivary gland invasion and transmission to the mammal, while ookinete motility and the infection of mosquitoes is not yet affected. Strikingly, the efficiency of mosquito colonization as measured by the presence of oocysts was not affected by specifically tuning the force generation capacity of myosin. Instead, a similar hierarchy appears with myosin mutants lacking some force generation capacity still being able to transmit efficiently, while the S19A mutant showed the largest defects in sporozoite migration and salivary gland colonization. This strongly suggests that the actin-myosin motor machinery was shaped and tuned to highest efficiency by the need of the sporozoites for their long and elaborate journey from the oocysts in the mosquito midgut to the hepatocyte in a mammalian liver. Strikingly, of all the barriers the entry into the salivary gland appears as the most formidable.

On a molecular engineering note: we first sought to use our established mutagenesis approach to alter *myoA* but found that the necessary change of the endogenous 3’UTR for the 3’UTR of *dhfs* led to decreased expression of *myoA* in early oocysts, which impacted the capacity of the sporozoites to enter salivary glands. A similar approach for generation of MyoA-GFP readily yielded salivary gland invasion (Green *et al*, 2017). However, the produced line was not clonal and the presence of wt *myoA* in the oocysts might have produced enough MyoA to compensate for the loss due to the unnatural 3’UTR. We previously used a gene-in-marker-out approach to mutate actin, which was also based on the generation of an initial recipient line where the 3’UTR of actin 1 was replaced by the 3’UTR of *dhfs* to facilitate gene replacement (Douglas *et al*, 2018). While this parasite line acted as a recipient for further modifications in the blood stages and was thus not required for mosquito infection, a strongly reduced salivary gland entry by this modified line was observed as well, suggesting that expression of not only *myoA* but also *actin 1* is affected by a modified 3’UTR (Ross Douglas, unpublished). To avoid similar problems with other proteins it might be advisable to use a gene-in-marker-out approach that leaves the 3’UTRs unaffected or a CRISPR/Cas9 system for subtle mutagenesis (Shinzawa *et al*, 2020).

Finally, we noted that it was possible to replace the *P. berghei myoA* gene for that of *myoA* from *P. falciparum*. However, the resulting parasites grew much slower in the blood and had deficits in infecting the mosquito. Multiple explanations are possible for these observations. Firstly, the myoA gene from *P. falciparum* was codon modified and hence the gene might not have been transcribed in sufficient amounts, as we noted recently for the gene encoding alpha-tubulin 1 (Spreng *et al*, 2019). Alternatively, the subtle differences in the amino acid sequence of the two myosins might be enough to cause the phenotype. Due to the additional problems associated with the 3’UTR we chose not to follow-up on these differences.

In conclusion, the presented data suggests that sporozoites need S19 phosphorylation to efficiently enter mosquito salivary glands and infect the mammalian host, consistent with the observation that MyoA is phosphorylated in sporozoites (Swearingen *et al*, 2017). Entry into salivary glands emerges as the strongest barrier in parasite transmission which the parasites need to overcome and represents the defining challenge in the parasite life cycle. This likely shaped the evolution of the motility and invasion machinery of *Plasmodium* and might constitute a similar barrier for other parasites that are transmitted from the salivary glands through the bite of an arthropod.

## Materials and Methods

### Ethics statement

Animal experiments were performed according to FELASA and GV-SOLAS guidelines and approved by the responsible German authorities (Regierungspräsidium Karlsruhe). *Plasmodium* parasites were maintained in NMRI or Swiss mice, transmission experiments were carried out using C57Bl/6 mice. Mice were obtained from JANVIER or Charles River Laboratories.

### Generation of mutant parasite lines

Cloning was performed using restriction enzymes or Gibson assembly. PCR fragments were amplified from genomic or plasmid DNA using the primers listed in **Supplementary Table 1**. Transfection vectors for parasite lines PbMyoA 3’dhfs, PbMyoA-GFP 3’dhfs and PfMyoA 3’dhfs were generated from vector Pb262 (Klug & Frischknecht, 2017). The myoA 5’UTR and 3’UTR were cloned into this vector to enable double crossover homologous recombination. GFP was fused to MyoA via a 4-amino acid alanine linker. A codon modified version of PfMyoA, kindly provided by Jake Baum (Blake *et al*, 2020), was cloned into the transfection vector to generate parasite line PfMyoA 3’dhfs.

For the introduction of point mutations into MyoA, a recipient line was produced from a p32 vector kindly provided by Jessica Kehrer. The myoA 5’UTR and open reading frame (ORF) were cloned into this vector to enable double crossover homologous recombination. The 5’UTR of ama1 was amplified and cloned into the vector. For the generation of parasite lines MyoA E6R, MyoA S19A and MyoA S19D, *myoA* was amplified with primers that were designed to introduce the mutations and cloned into vector Pb262.

The transfection vectors were linearized via restriction digest with KpnI or BamHI and SacII before transfection. *Plasmodium berghei* ANKA parasites were used for genetic modifications. Transfection and generation of isogenic parasite lines was carried out essentially as described before (Janse *et al*, 2006; Klug & Frischknecht, 2017). DNA was prepared using ethanol precipitation and electroporation of purified schizonts was carried out using Nucleofactor technology (Lonza). Transfection mixtures were then injected intravenously into a mouse. Transfected parasites were positively selected with 0.07 mg/mL pyrimethamine or negatively selected with 1.5 mg/ml 5-FC added into the drinking water of the mice. Clonal parasite lines were obtained by limiting dilution. Successful mutagenesis was verified via genotyping PCR with the primers listed in **Supplementary Table 2** and point mutations were additionally verified by sequencing of the modified locus.

### Mosquito infection and sporozoite isolation

*Anopheles stephensi* mosquitoes were infected and sporozoites isolated from midguts, hemolymph and salivary glands as described previously (Klug & Frischknecht, 2017). Briefly, for mosquito infections, mice were infected by intraperitoneal injection of frozen stocks (150–200 ml). After 3–5 days, the infected mice were checked for presence of gametocytes by placing a drop of tail blood on a microscope slide followed by incubation at room temperature for 10–12 min. During this time, flagellated gametes mature and exit in a process termed exflagellation. If 1–2 exflagellation events per field were observed at 40x magnification, mice were anesthetized and fed to mosquitoes. Experiments with hemolymph and salivary gland sporozoites were performed 13 to 16 and 17 to 25 days post mosquito infection, respectively. Oocysts were counted from dissected midguts following mercurochrome staining: Midguts were permeabilized for 20 min with 1% Nonidet P40 in PBS, stained with 0.1% mercurochrome in PBS for 30 min, mounted in a small amount of PBS, covered with a coverslip, and counted using a 10× objective on a Zeiss CellObserver microscope. Midguts and salivary glands were dissected with a pair of needles in PBS and placed on ice until further use. Sporozoites were released by gently crushing the collected organs with a plastic pestle. Sporozoites were isolated from the hemolymph by cutting the last segment of the abdomen with a syringe and flushing with RPMI (supplemented with 50,000 units/L penicillin and 50 mg/L streptomycin) by inserting a long-drawn Pasteur pipette into the lateral side of the thorax. The hemolymph was drained from the abdomen, collected on a plastic foil and transferred to a reaction tube. Sporozoites were counted using a Neubauer chamber and total numbers extrapolated according to the dilution.

### Cell migration assays

Cell migration assays on glass or in polyacrylamide hydrogels were performed and analysed as described before (Douglas *et al*, 2018; Ripp *et al*, 2021). Briefly, isolated sporozoites in RPMI medium supplemented with 50,000 units/l penicillin, 50 mg/l streptomycin and 3% bovine serum albumin (BSA) were pipetted into a 96-well optical bottom plate (Nunc) and centrifuged at 1,000 rpm for 3 min. For 3D hydrogel assays, whole infected salivary glands dissected into 30 µl of medium were sandwiched between a glass coverslip (22 × 22 mm) placed on top of a microscope slide and a 3% AA/0.03% BIS hydrogel. Imaging was performed at room temperature on an inverted Zeiss CellObserver microscope. Images were recorded every second for hemolymph sporozoites or every three seconds for salivary gland sporozoites for a total time of three minutes. Sporozoite speed was determined by the manual tracking plugin from FIJI (Schindelin *et al*, 2012).

### Force measurements

Force experiments were performed on a Nikon Eclipse TI microscope equipped with an MMI CellManipulator with a 1070 nm laser (8W). Sample preparation was carried out as described earlier (Quadt *et al*, 2016). In short, infected salivary glands of 3-5 mosquitoes dissected in 35 µl of medium were smashed, flown into a self-made flow chamber and incubated for 10 minutes. If sporozoites were attached and motile, the chamber was washed with streptavidine-coated polystyrene beads (1.5 - 1.9 µm, Kisker Biotech) in 3% BSA/RPMI. Laser trap calibration was performed with the MMI Cell Tool software by adjusting the escape force of the bead to about 30 pN. The escape force was determined by harmonic bead oscillation at the z-focus of later experiments according to the Stoke’s equation with the known viscosity of the medium and bead diameter. A bead was captured at a defined laser power of 30 pN and placed onto the apical end of a moving sporozoite. The percentage of motile sporozoites that could pull the bead out of the trap towards the rear end of the sporozoite was quantified. Experiments were performed at room temperature.

### qRT-PCR

To determine expression levels of MyoA in wt and PbMyoA 3’dhfs parasites, total RNA of 15-20 mosquito midguts (day 10 post-infection) and purified schizonts (10 million parasites) was isolated with Qiazol reagent according to the manufacturer’s protocol (Invitrogen). RNA was purified with the Direct-zol RNA MicroPrep kit (Zymo Research) and cDNA synthesis was generated using the First-Strand cDNA synthesis kit (Thermo Fisher Scientific) according to the manufacturer’s protocols. The quantitative PCR was performed using SYBR Green PCR Master Mix (Life Technologies) including ROX dye and was measured with the Bio-Rad CFX96 Real-Time System. The 18S rRNA gene was used as a reference, and across-run differences were normalized using a calibrator sample. Relative copy numbers were calculated by applying the ΔΔCt methodology. The sequences of the gene-specific primers used are shown in **Supplementary Table 3**.

### Analysis

Graphs were created using R. Statistical analysis was performed in R. Figures were generated using Inkscape.

## Acknowledgements

We thank Miriam Reinig for rearing *Anopheles stephensi* mosquitoes, Joachim Spatz for access to and Katharina Quadt for set-up, calibration and introduction of the new laser tweezers as well as Jake Baum, Ross Douglas, Franziska Hentzschel and Anne Houdousse for helpful discussions and comments on the manuscript. This work was funded by grants from the Human Frontier Science Program (RGY0066/2016) and the Deutsche Forschungsgemeinschaft (DFG, German Research Foundation, DFG project number 240245660 – SFB 1129). FF is a member of CellNetworks Cluster of excellence at Heidelberg University and SFB 1129. JR was a member of Heidelberg International Graduate School for the Biosciences (HBIGS) and XS and MTN members of the Master Program for Molecular Biotechnology at Heidelberg University. We acknowledge the essential microscopy support from the Infectious Diseases Imaging Platform (IDIP) at the Center for Integrative Infectious Disease Research.

## Supplementary Figures and Tables

**Supplementary Figure 1.**
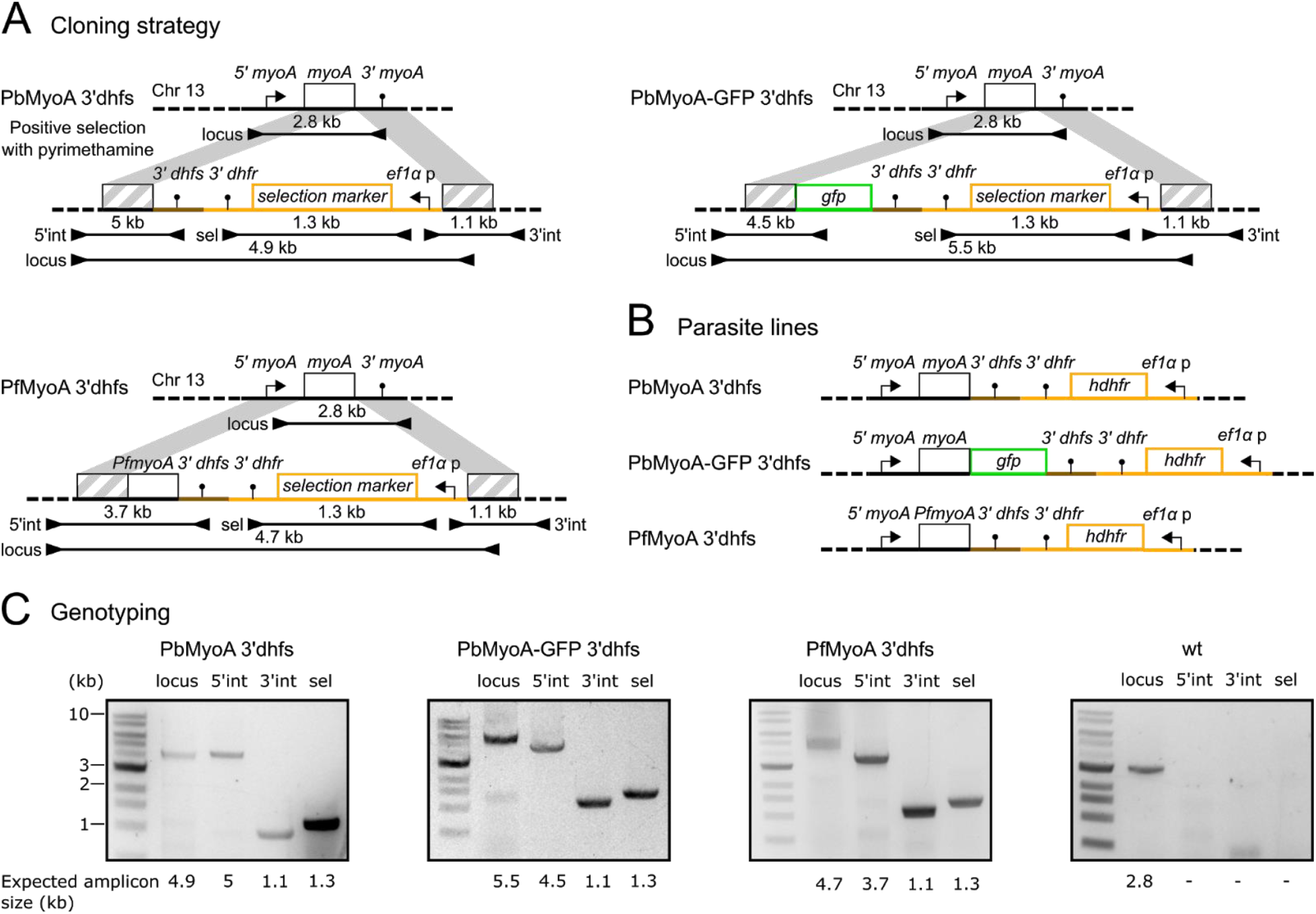
Generation of PbMyoA 3’dhfs, PbMyoA-GFP 3’dhfs and PfMyoA 3’dhfs parasite lines. (A) Scheme of cloning strategy. Homology regions are indicated in grey. Shown below are the amplified PCR fragments to test for successful integration, the primer binding site and the amplicon size. All parasite lines were obtained by positive selection using pyrimethamine. Note that the scheme is not drawn to scale. (B) MyoA locus of all parasite lines that were generated using the cloning strategy depicted in (A). (C) Correct integration was verified via genotyping PCR. Given below are the expected sizes of the amplicons.

**Supplementary Figure 2.**
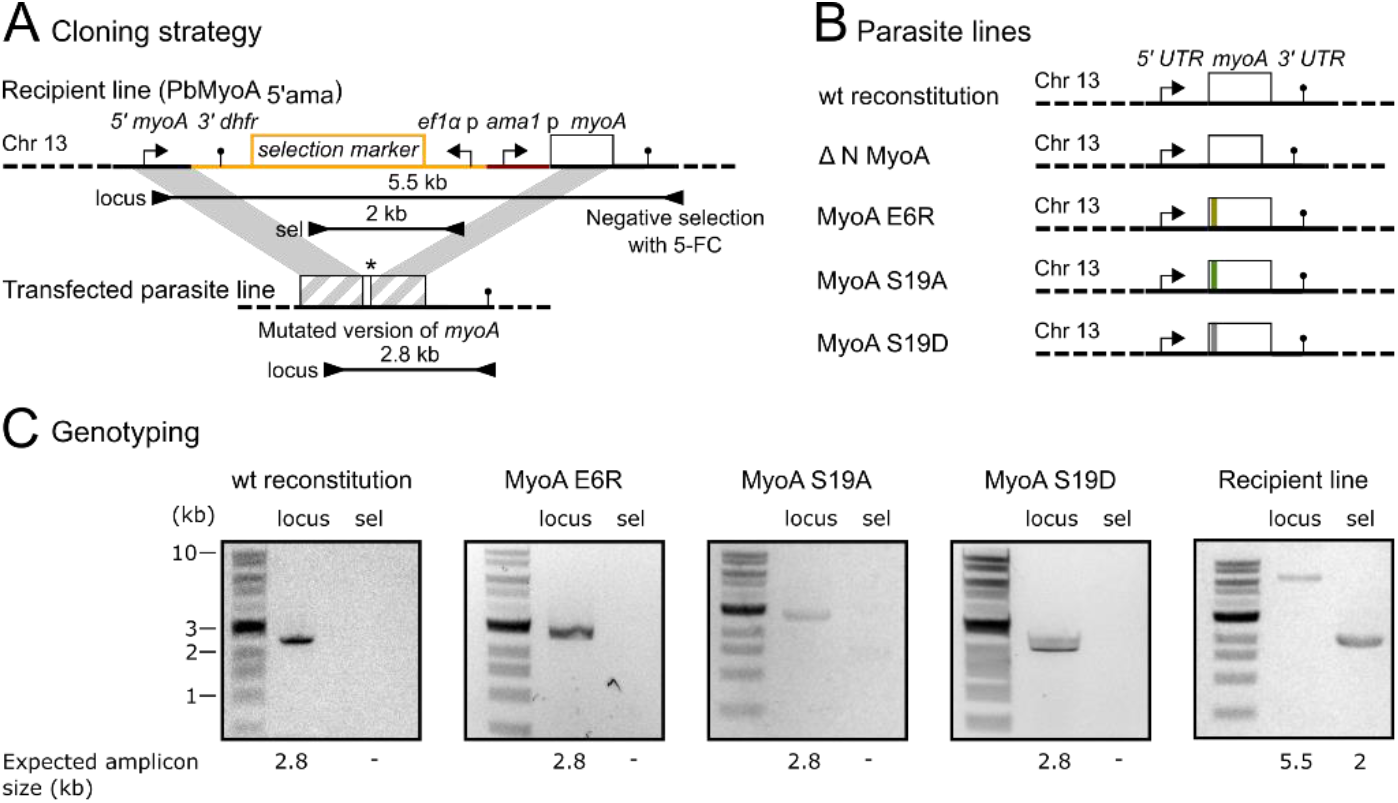
Generation of point mutants. (A) Scheme of cloning strategy. Homology regions are indicated in grey. Shown below are the amplified PCR fragments to test for successful integration, the primer binding site and the amplicon size. All parasite lines were obtained by negative selection using 5-FC. Note that the scheme is not drawn to scale. (B) MyoA locus of all parasite lines that were generated using the cloning strategy depicted in (A). (C) Correct integration was verified via genotyping PCR. Given below are the expected sizes of the amplicons.

**Supplementary Table 1.**
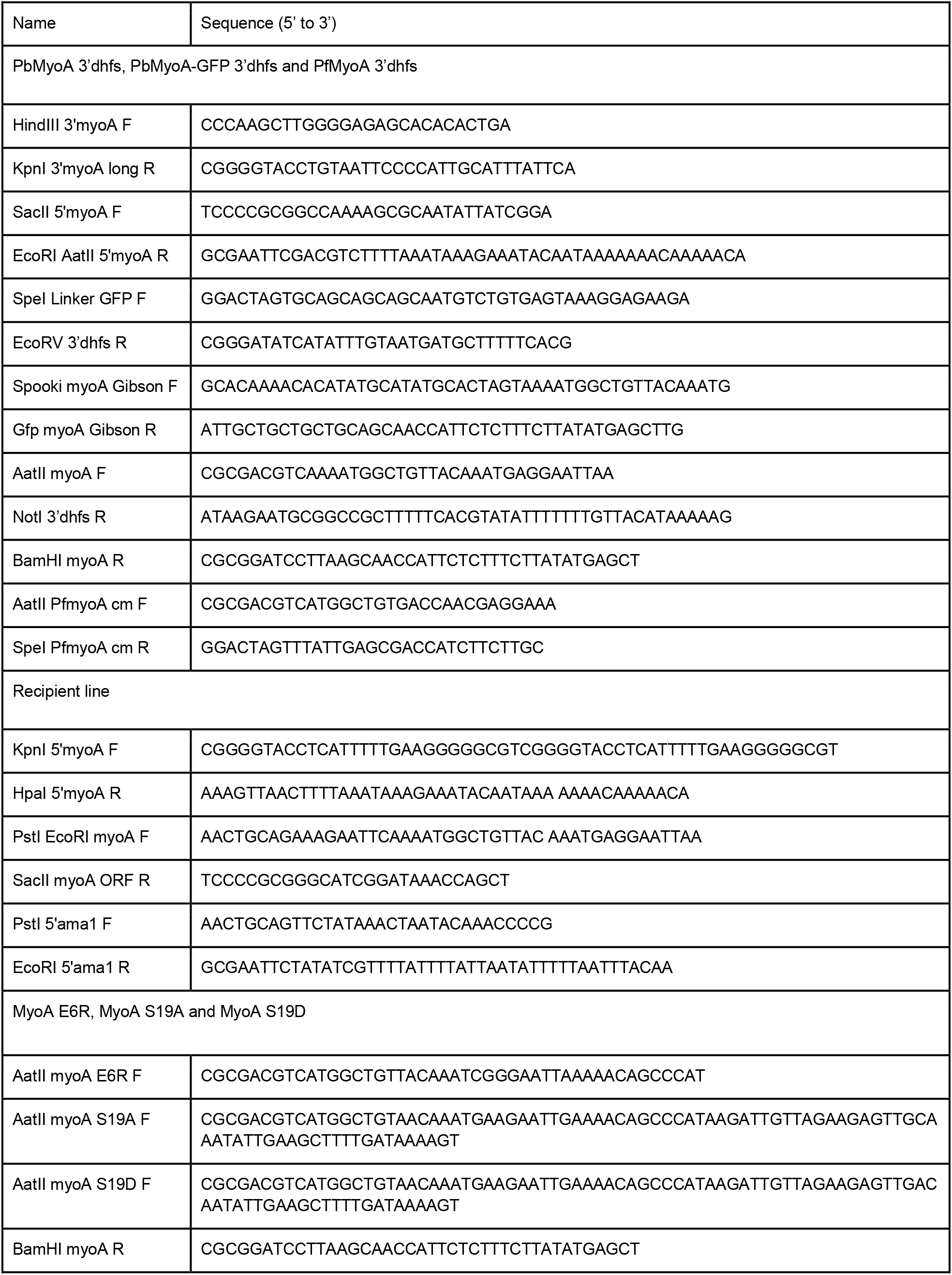
(Primers used for cloning of transfection vectors)

**Supplementary Table 2.**
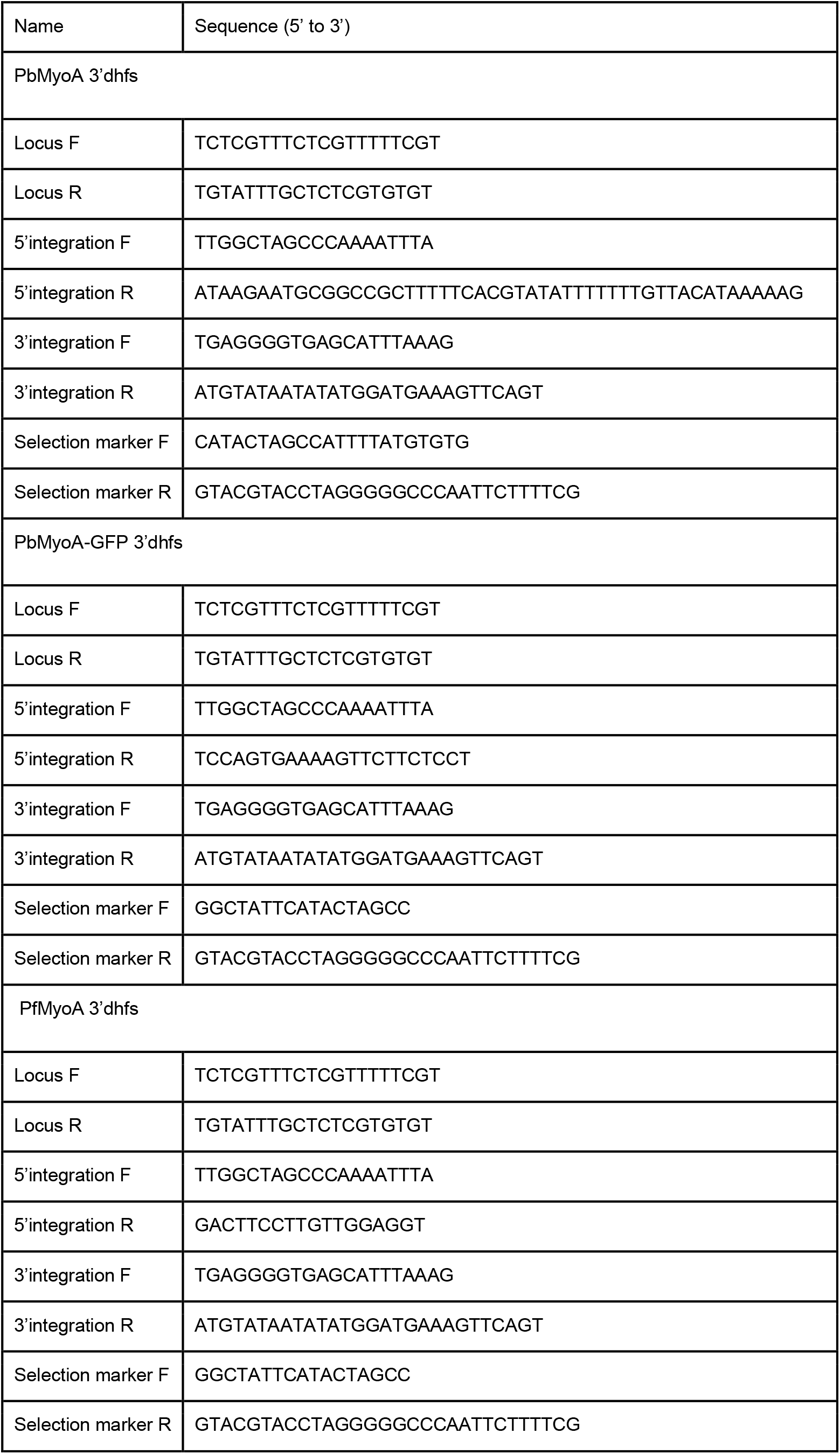
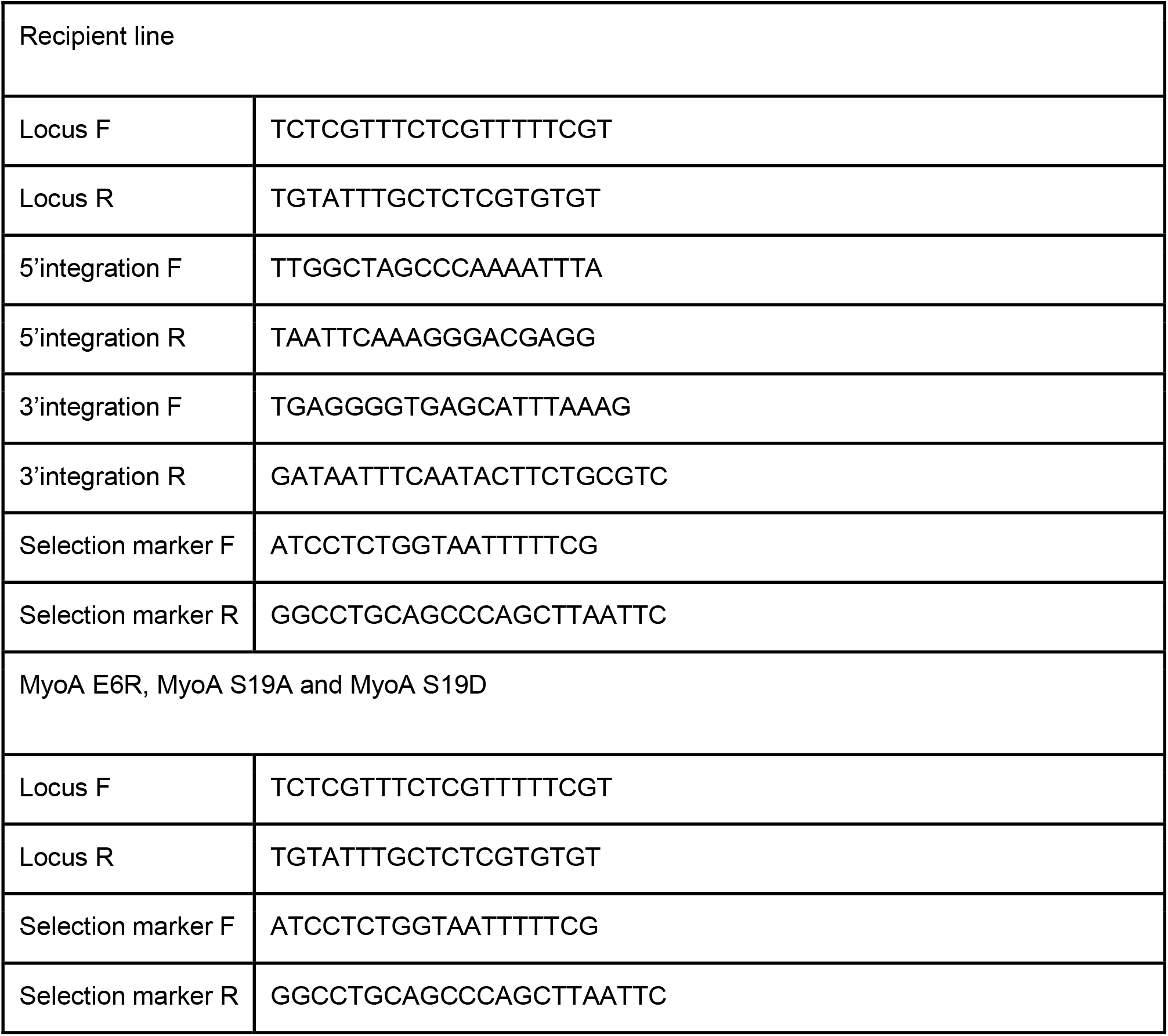
(Genotyping Primers)

**Supplementary Table 3.**
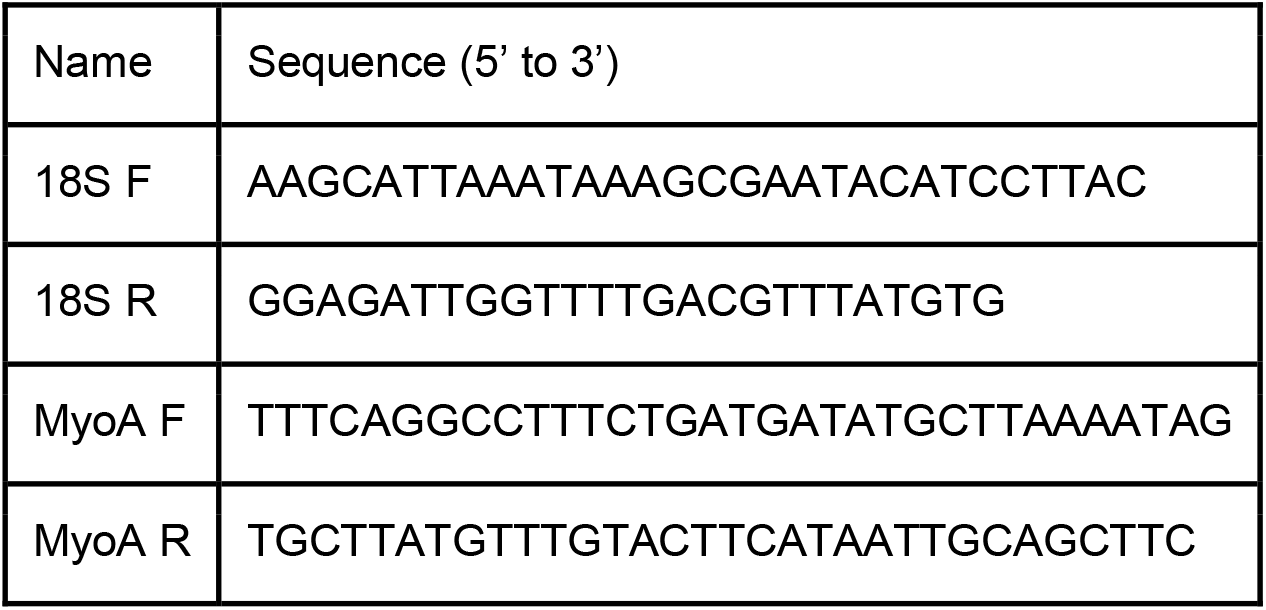
(Primers for qRT-PCR)

